# ANYI: The ANnotated Yeast Interactome

**DOI:** 10.64898/2026.04.30.721908

**Authors:** Daniel A. Nissley, Muskan Goel, Xavier Castellanos-Girouard, Charles P. Kuntz, Yiqing Wang, M. Shahid Mukhtar, Adrian Serohijos, Jonathan P. Schlebach

## Abstract

Although several existing protein–protein interaction (PPI) databases provide yeast PPI data, none unify large-scale network topology information with detailed biophysical, proteostasis, and regulatory annotations in a single protein-centric framework. To address this gap, we developed the ANnotated Yeast Interactome (ANYI), an open, integrated resource that combines experimental yeast PPIs with sixteen feature annotation types, including protein abundance, half-life, disorder content, post-translational modifications, conformational stability, chaperone interactions, sequence, and structure. ANYI integrates 3,927 proteins with 155 annotation features, forming a unified matrix that enables systematic cross-layer analyses. Available via GitHub and Docker Hub with an interactive network browser for broad accessibility, ANYI provides both experienced and beginner computational scientists with tools to investigate the yeast interactome. For example, users can directly test whether highly connected hub proteins exhibit distinct stability, disorder, or proteostasis signatures relative to peripheral nodes.

**AVAILABILITY AND IMPLEMENTATION:** The code used to assemble ANYI is available on GitHub at https://github.com/NCEMS/energetic-origins-of-PPI-connectivity and the database itself and interactive browser tool are available on Docker Hub as dannissleypsu/anyi-browser:v1.0.2.

## 1 INTRODUCTION

Cells rely on networks of protein-protein interactions (PPIs) to coordinate metabolism and signaling. Efforts to map these networks and rationalize their collective behavior emerged in the 1990s and early 2000s with the introduction of the yeast two-hybrid screening method (Fields and Song 1989), which was later developed into a high-throughput technique suitable for proteome-wide analyses (Fromont-Racine, Rain, and Legrain 1997). In the early 2000s, several comprehensive yeast two-hybrid studies profiled thousands of interactions between proteins in yeast (Uetz *et al*. 2000, Ito *et al*. 2001), providing the first high-coverage yeast interactomes. Around the same time, advances in mass spectrometry began to enable identification of PPIs with techniques that do not rely on reporter gene expression to detect interactions (Gavin *et al*. 2002, Ho Y *et al*. 2002). These mass-spectrometry-based methods led eventually to the publication of The Yeast Interactome, collating 31,004 interactions between 3,927 yeast proteins, covering approximately two thirds of the yeast proteome (Michaelis *et al*. 2023).

Within the interactome view of protein function, a protein’s role is defined not only by its enzymatic activity or interaction with signaling molecules but also its position within the PPI network. Contextualizing various biological and biochemical data within a PPI network has led to the discovery of several emergent network properties. For example, Jeong and co-workers found that proteins that form highly connected hubs within PPI networks are three times as likely to be essential than proteins with a small number of interactions with other proteins (Jeong *et al*. 2001). Studies have also shown that highly connected proteins may tend to have higher copy numbers (Ivanic *et al*. 2009), undergo sequence evolution more slowly (Fraser, Wall, and Hirsh 2003), and are more likely to contain disordered regions (Haynes *et al*. 2006).

Motivated by the wealth of discoveries that have emerged from studying the yeast proteome, we present ANYI, the ANnotated Yeast Interactome. ANYI is built upon the high-coverage yeast interactome generated by the Mann Lab (Michaelis *et al*. 2023), which leveraged systematic, quantitative affinity-purification mass spectrometry with well-characterized detection limits, high reproducibility, and stringent statistical filters, resulting in a dense, high-confidence PPI network. ANYI associates 16 classes of experimental and computational information with proteins (network nodes) to enable comparisons and hypothesis testing within the context of the interactome. These annotations include predicted disorder content, protein abundance, entanglement status, and protein stability predictions from three methods. Importantly, ANYI does not alter the underlying PPI network but instead enriches each node with standardized, harmonized annotations to enable systems-level synthesis. An Open-Source container for ANYI enables the exploration of the yeast interactome by users with minimal computational science training.

## 2 METHODS & RESULTS

### 2.1 Annotating the yeast interactome

ANYI integrates 16 complementary experimental and computational data classes into a unified, protein-centric annotation framework using Saccharomyces Genome Database (SGD) systematic names as unique identifiers (Engel *et al*. 2025). Data integration was carried out using Python code in combination with Open Source and licensed computational tools. Starting from the Mann Lab’s PPI network (Figure 1A), we computed network centrality features (*e*.*g*., degree and betweenness centrality) and incorporated primary sequence information from the SGD and sequence-derived predictions (Hallgren *et al*. 2022, Teufel *et al*. 2022) alongside curated UniProt annotations (Bateman *et al*. 2025). Structural features including predicted disorder content (Lotthammer *et al*. 2024), protein stability predictions (Ghosh and Dill 2010, Alford *et al*. 2017, Cagiada, Ovchinnikov, and Lindorff-Larsen 2025), domain annotations (Blum *et al*. 2025), and entanglement status (Rana *et al*. 2024) were added to capture intrinsic protein biophysical features. Context within the broader proteome is incorporated in form of half-life (Christiano *et al*. 2014, Martin-Perez and Villén 2017), steady-state expression level (Ho B, Baryshnikova, and Brown 2018), translation efficiency (Fang *et al*. 2018), experimental and predicted post-translational modifications (Deutsch *et al*. 2023, Shrestha *et al*. 2024), chaperone interactions (Rizzolo *et al*. 2017), oligomerization state (Balu *et al*. 2025), and thermal stability information (Jarzab *et al*. 2020). Finally, complementary yeast two-hybrid network data is included as an independent PPI modality (Yu *et al*. 2008). At each integration step, datasets were standardized to common formats and merged via explicit key-based mapping to produce a single consolidated annotation table. All quantitative features are provided on their native scales to allow flexible downstream normalization depending on the intended context. The result is an approximately 600,000-dimensional matrix (3,927 nodes x 155 annotations, Figure 1B) of annotations. All annotations are summarized in Table S1, and data sources and processing steps are described in detail in the Supplementary Methods.

**Figure 1.**
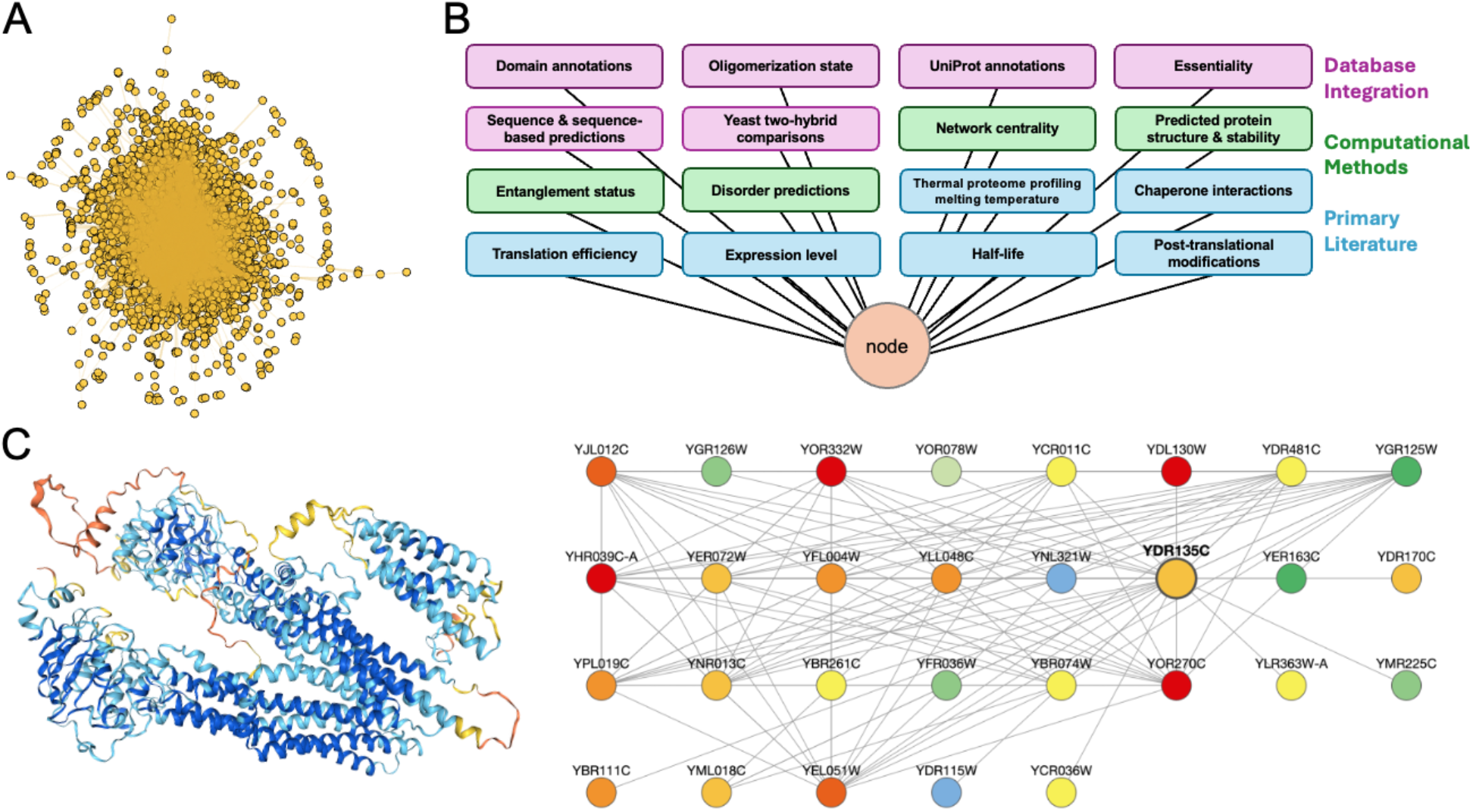
(A) The Yeast Interactome from the Mann Lab, which contains 3,927 nodes and 31,004 edges, is the basis for ANYI. (B) Each node in ANYI is associated with 16 classes of information derived from public databases, computational predictions, and datasets from the primary literature. (C) The ANYI Browser allows nodes within the Yeast Interactome to be viewed in their network and biophysical context. Here, the portions of the viewer that display the AlphaFold2 (Jumper *et al*. 2021) structure of YDR135C (left) and its ego network with nodes colored by molecular abundance (right) are shown.

### 2.2 Testing hypotheses with ANYI

Generating and testing hypotheses with ANYI can be accomplished in two ways: First, users can load the database into Python using Pandas and utilize the full suite of Python libraries for data analysis and statistics. We demonstrate how to use ANYI to test the association between essentiality and node degree (degree centrality) with Fisher’s Exact Test in the examples folder of the GitHub. Beyond this classical example, ANYI enables multi-layer analyses such as testing whether hub proteins are enriched for intrinsic disorder, whether chaperone-interacting proteins exhibit lower predicted folding stability, or whether entangled proteins occupy distinct topological positions within the network. Second, users can employ a Jupyter Notebook-based graphical user interface, the ANYI Browser, to explore the diverse connections between yeast proteins. This node-centric search and exploration tool enables users to query ANYI using SGD systematic names to view an instant readout on key quantities and structural information (Figure 1C, left). A customizable display of the ego network for the selected protein enables figure generation and contextualization of biophysical and biochemical information (Figure 1C, right). Unlike general-purpose visualization platforms, the ANYI Browser is pre-configured to couple structural, stability, and expression annotations directly to network context without requiring manual data import or attribute mapping. Both data analysis methods are available via the ANYI container on DockerHub.

## 3 DISCUSSION

Though many data modalities from both experimental and computational methods are available in the molecular and cellular sciences, data integration remains a significant barrier to data synthesis research. Even in mature fields, available data are fragmented across multiple federated databases and portals that emphasize different layers of information. For example, Reactome focuses on manually curated human PPIs (Milacic *et al*. 2024); STRING aggregates interactions from many resources to compute a single score (Szklarczyk *et al*. 2023); BioGrid prioritizes relationships across layers including chemical interactions to facilitate drug discovery (Oughtred *et al*. 2021). None of these resources provide an integrated, yeast-specific framework that simultaneously exposes network topology, quantitative proteome measurements, structural stability metrics, disorder predictions, and proteostasis annotations in a single harmonized dataset.

Investigators who want to ask integrative questions - linking network topology to protein biophysics, proteostasis, and regulation - typically spend substantial effort on the practical aspects of data integration. Tasks like resolving redundant, conflicting, or context-specific annotations, or tracking code provenance and versions, often require highly specific knowledge. These practical requirements become acute when the desired analysis includes the implementation of cutting-edge computational workflows, which may be incompletely documented and difficult for non-experts to treat as anything other than a black box. Many potentially informative cross-dataset analyses are thus either not attempted or are executed in an *ad hoc* manner that is difficult to reproduce and extend, limiting the insights gained from existing data.

By integrating diverse experimental and computational data into the context of The Yeast Interactome, ANYI lowers these barriers by providing an analysis-ready, protein-centric environment for systematic hypothesis generation and testing in *S. cerevisiae*. ANYI enables questions such as whether network hubs are biophysically distinct, whether proteostasis constraints correlate with topological centrality, or whether specific structural classes preferentially occupy certain network neighborhoods. Rather than functioning as another primary repository, ANYI is explicitly structured as a synthesis layer: it harmonizes identifiers, standardizes and annotates features, and packages them in a form that can be directly queried, filtered, and modeled without extensive reprocessing. This design enables users to move quickly from conceptual questions to quantitative analyses. ANYI shifts effort away from one-off data wrangling and toward interpretable, reproducible synthesis to enable downstream analysis and data reuse. We note that ANYI inherits limitations from the underlying interactome and annotation datasets, including reliance on a static PPI network, incomplete proteome coverage, and potential biases in experimental measurements. Future updates may incorporate condition-specific interaction data and additional regulatory layers as they become available.

## 4 CODE AVAILABILITY

All required code is available on GitHub at github.com/NCEMS/energetic-origins-of-PPI-connectivity. Full reproduction of some pipeline steps requires additional software packages (*e*.*g*., Rosetta, SignalP) that can be downloaded free of charge for academic users. See the README on GitHub for details.

## 5 DATA AVAILABILITY

A Docker container available on Docker Hub as dannissleypsu/anyi-browser:v1.0.2 provides the ANYI Browser environment to users on most systems (see the GitHub for download and usage instructions). All processed and raw data are available on the CyVerse data store (see GitHub for details); a free CyVerse account is required for access.

## 6 ACKNOWLEDGEMENTS

We thank Eugene I. Shakhnovich for his scientific input. This work was supported by the National Science Foundation (NSF) through the NSF National Synthesis Center for Emergence in the Molecular and Cellular Sciences (NCEMS) under Grant NSF MCB-2335029. A.W.R.S. acknowledges support from the Canadian Institute of Health Research grant MOP-G-408523, Natural Sciences and Engineering Research Council of Canada grant RGPIN-2016-06566, and the Canada Research Chairs. J.P.S acknowledges support from the National Institute of General Medical Sciences award R35GM152086. M.S.M. acknowledges support from National Science Foundation award IOS-2038872, and OIA-2418230. X.C.G. acknowledges support from the Natural Sciences and Engineering Research Council of Canada – Masters Canada Graduate Scholarship and Fonds de Recherche du Québec en Santé – Masters Training Scholarship.

