## Supplementary methods & data for "ANYI: The ANnotated Yeast Interactome"

The 16 sections below correspond to the 16 annotation classes within ANYI. Each methods section corresponds to a set of rows in Table S1 that define the column identifiers in ANYI that can be used to access protein-level information for that annotation class.

To interact with the ANYI database itself, follow the instructions on GitHub to download and install the ANYI Browser Docker container, which includes an example of loading ANYI into Python for analysis.

**1 - Network centrality calculations.** The Cytoscape (Shannon *et al.* 2003) session file for The Yeast Interactome was downloaded from www.yeast-interactome.org (Michaelis *et al.* 2023) and used as the basis for downstream annotations. This session file was opened in Cytoscape v3.10.3, weighted k-shell decomposition values (Wei *et al.* 2015) calculated with the wk-shell-decomposition app, and the node and edge tables exported. Network models were constructed using these node and edge tables in Python with NetworkX (Hagberg, Schult, and Swart 2008) v3.4.2 and its internal functions used to compute degree centrality, betweenness centrality, eigenvector centrality, closeness centrality, load centrality, and PageRank (with damping coefficient 0.85) for the full network (3,927 nodes). NetworkX was also used to compute information centrality using the largest connected subgraph (3,839 nodes).

**2 - Sequence information and sequence-based predictions.** Protein sequences (Genome Release 64-5-1) were retrieved from the SGD for the S288C reference genome and associated with network nodes based on their SGD open reading frame identifiers. DeepTMHMM (Hallgren *et al.* 2022) v1.0.24 was used to classify proteins as globular or transmembrane, and SignalP 6.0h fast (Teufel *et al.* 2022) was used to predict signal peptide cleavage sites. Predicted signal peptides were removed based on these cleavage site predictions, and all subsequent analyses used the trimmed sequences.

**3 - UniProt annotation.** UniProt accession numbers were assigned to each node using the "idmapping" file for S288C yeast from the UniProt FTP site ftp.uniprot.org. Each node was then annotated with information from UniProt (Bateman *et al.* 2025), including post-translational modifications, subcellular localization, Gene Ontology (GO) terms, and protein function information. Gene ontology terms were also converted to a human-readable format using obolibrary (Ashburner *et al.* 2000) and the go-basic.obo reference file (Aleksander et al. 2026). UniProt annotations were extracted from a copy of uniprot_sprot.xml downloaded from the UniProt FTP site ftp.uniprot.org. Copies of all source files are stored on CyVerse (see GitHub).

**4 - Intrinsic disorder content and property predictions.** Intrinsically disordered regions (IDRs) were predicted from protein sequences using metapredict v3.0 (Lotthammer, Hernández-García *et al.* 2024). Sequence and ensemble properties were computed for all IDRs with CIDER (Holehouse *et al.* 2017) and ALBATROSS (Lotthammer, Ginell *et al.* 2024) functionality within sparrow from github.com/idptools. The fraction of disorder predicted within each protein was compared to various percentile thresholds computed from the disorder content for all entries for S288C proteins in the DisProt database (Nugnes *et al.* 2026).

**5 - Protein stability prediction.** Folding free energies were estimated using two approaches for proteins predicted by DeepTMHMM to not be transmembrane proteins: (1) Equation 1 from Ghosh & Dill (2010) (Ghosh and Dill 2010), which depends only on protein length and system temperature (set to 300 K), & (2) the ESM-IF-based model from Cagiada et al. (Cagiada, Ovchinnikov, and Lindorff-Larsen 2025), which requires AlphaFold2-predicted structures (Jumper *et al.* 2021). For this latter calculation, AlphaFold2 structure predictions (version 4) for the yeast proteome were obtained from the European Bioinformatics Institute (EBI) (Varadi *et al.* 2024). For proteins with predicted signal peptides or a mismatch between the sequences of the EBI structure and the SGD reference sequences, new structure predictions were carried out with AlphaFold2. All structures were also scored using Rosetta release 371 with the ref2015 scoring function (Alford et al., 2017). Prior to Rosetta scoring, AlphaFold2 structures were relaxed with the FastRelax protocol in Rosetta using ten independent replicas per protein. Python code to implement the Cagiada et al. 2025 model is based on the example Jupyter Notebook from https://github.com/KULL-Centre/_2024_cagiada_stability.

**6 - Protein half-life data integration.** Protein half-life information from two large-scale mass spectrometry studies (Christiano *et al.* 2014, Martin-Perez and Villén 2017) was incorporated using simple identifier-based joins. The data from Christiano *et al.* 2014 and Martin-Perez and Villén 2017 were obtained from Table S1 and Data S1, respectively, of the corresponding publications.

**7 - Protein expression level data integration.** Protein expression levels from a meta study (Ho, Baryshnikova, and Brown 2018) were incorporated using an identifier-based mapping. Specifically, the data from Ho et al. 2018 Table S4, representing the molecules per cell data after correction for autofluorescence, were integrated into ANYI.

**8 - Translation efficiency data integration.** Pre-computed translation efficiency values for yeast from the sckit-ribo package (Fang *et al.* 2018) were integrated into ANYI. The source file, skr_weinberg_genesTE.csv, is available on the scikit-ribo_manuscript GitHub in the blob/master/Data directory.

**9 - Predicting post-translational modifications.** In addition to the curated PTM information included from UniProt annotations (see above), yeast phosphorylation sites from PTMeXchange mass spectrometry data (PXD071918) downloaded from ProteomeXchange (Deutsch *et al.* 2023) were associated with each network node. In addition, the locations of 19 classes of PTMs were predicted with the PTMGPT2 language model (Shrestha *et al.* 2024).

**10 - Entanglement status data integration.** Binary entanglement status information computed from both AlphaFold2 and experimental structures was included for each protein based on results from Rana & Sitarik *et al.* 2024. Here, entanglements are defined as non-covalent lasso structures (Rana *et al.* 2024).

**11 - Chaperone interaction data integration.** The list of chaperones and co-chaperones in yeast from Rizzolo *et al.* 2017 Table S1 was used to define a list of chaperone nodes (Rizzolo *et al.* 2017). All nodes that share an edge with a chaperone node in The Yeast Interactome are considered to interact with that chaperone.

**12 - Oligomerization state data integration.** The Complex Portal (Balu *et al.* 2025) reference file for yeast, 59292.tsv, was downloaded from EBI and parsed to extract information at the individual protein level. Within ANYI, various binary columns encode if each protein participates in the formation of specific types of complexes (*e.g.*, homodimer, homotrimer, heterodimer) or in the formation of complexes with unknown stoichiometry.

**13 - Domain annotations**. The InterPro (Blum *et al.* 2025) database file protein2ipr.dat.gz was downloaded from EBI and parsed to extract domain annotations for each protein in ANYI (see Table S1 for a summary of all domain annotation methods included). In addition to a list of the domains identified by each method, we also include a summary of the number of domains identified for each protein by each method.

**14 - Essentiality data integration.** The SGD web browser was used to select all S288C proteins with a phenotype of “inviable”, which indicates proteins that are essential to growth, and these annotations then mapped to network nodes.

**15 - Yeast two-hybrid PPI data integration.** Data were obtained from the CCSB Interactome Database (Yu *et al.* 2008) and network centrality calculations carried out as described for The Yeast Interactome data in section **1 - Network centrality calculations**. The network centralities within the yeast two-hybrid interactome were associated with each node within the mass spectrometry-derived Yeast Interactome from the Mann Lab to provide a secondary set of reference centralities.

**16 - Thermal proteome profiling data integration.** Proteome-wide thermal stability information was incorporated from the Meltome Atlas (Jarzab *et al.* 2020). Coefficient of determination values for melting curve fit line evaluation are reproduced in ANYI to enable the filtering of results by quality.

### Supplementary Table 1 – Annotations within ANYI

| Annotation Class | Index | Data Column Name | Explanation |
|---|---|---|---|
| 0 - Identifiers | 1 | node | SGD systematic name; primary identifier |
| | 2 | UniProtKB-AC | UniProt knowledge base primary accession; secondary identifier |
| 1 - Network centrality | 3 | _wkshell | Weighted *k*-shell decomposition value for this node |
| | 4 | degree_centrality | Degree centrality of this node |
| | 5 | betweenness_centrality | Betweenness centrality of this node |
| | 6 | eigenvector_centrality | Eigenvector centrality of this node |
| | 7 | closeness_centrality | Closeness centrality of this node |
| | 8 | load_centrality | Load centrality of this node |
| | 9 | pagerank | PageRank of this node |
| | 10 | information_centrality | Information centrality of this node |
| 2 - Sequence information and sequence-based predictions | 11 | has_verified_sequence | Boolean True if this node could be associated with an SGD Open Reading Frame, otherwise False |
| | 12 | sequence | The single-letter amino acid sequence of this protein from SGD |
| | 13 | DeepTMHMM_trimmed_sequence | Trimmed single-letter amino acid sequence resulting from removal of residues identified by DeepTMHMM to be part of signal sequence (S in output mask) |
| | 14 | DeepTMHMM_class | One from the list of GLOB (globular), TM (transmembrane, alpha helical), SP (has signal sequence), BETA (transmembrane, beta barrel), or SP+TM (alpha helical membrane protein with signal sequence) |
| | 15 | cleavage_site_start | Predicted start site of signal sequence cleavage from SignalP; if this value is i, cleavage is between i and i + 1 |
| | 16 | signalP_trimmed_sequence | Sequence predicted by SignalP to remain after cleavage of the signal peptide; if there was no signal sequence, this is identical to sequence |
| 3 - UniProt annotations | 17 | ProteinName | Primary name of the protein for this node from UniProt |
| | 18 | GO_terms | GO terms for this protein from UniProt |
| | 19 | GO_term_human_readable | GO terms for this protein from UniProt with their human-readable name included |
| | 20 | localization_keywords | Localization information from UniProt (UniProt annotation) |
| | 21 | parsed_functions | Functional information from UniProt |
| | 22 | parsed_PTMs | Post-translation modifications from UniProt annotations |

### Supplementary Table 1 (cont.) – Annotations within ANYI

| Annotation Class | Index | Data Column Name | Explanation |
|---|---|---|---|
| 4 - Disorder predictions | 23 | IDR_count | Integer number of discrete IDRs within the protein using a 30-aa cutoff |
| | 24 | IDR_sequences | Dictionary containing all IDR sequences (not just those meeting the 30 aa cutoff); keys are integers, matches IDR_ranges. Computed with metapredict v3 on the signalP_trimmed_sequence |
| | 25 | IDR_ranges | Dictionary containing IDR residues ranges for all IDR sequences (keys are integers, matches IDR_sequences) |
| | 26 | N_aa_disordered | Integer number of disordered residues in the protein |
| | 27 | disorder_fraction | Fraction of disordered residues in the protein |
| | 28 | is_disordered_0.05 | 1 if disorder_fraction is ≥ the 5th percentile of disorder fraction within DisProt for S288C proteins, otherwise 0 |
| | 29 | is_disordered_0.10 | 1 if disorder_fraction is ≥ the 10th percentile of disorder fraction within DisProt for S288C proteins, otherwise 0 |
| | 30 | is_disordered_0.15 | 1 if disorder_fraction is ≥ the 15th percentile of disorder fraction within DisProt for S288C proteins, otherwise 0 |
| | 31 | is_disordered_0.20 | 1 if disorder_fraction is ≥ the 20th percentile of disorder fraction within DisProt for S288C proteins, otherwise 0 |
| | 32 | is_disordered_0.25 | 1 if disorder_fraction is ≥ the 25th percentile of disorder fraction within DisProt for S288C proteins, otherwise 0 |
| | 33 | is_disordered_0.30 | 1 if disorder_fraction is ≥ the 30th percentile of disorder fraction within DisProt for S288C proteins, otherwise 0 |
| | 34 | is_disordered_0.35 | 1 if disorder_fraction is ≥ the 35th percentile of disorder fraction within DisProt for S288C proteins, otherwise 0 |
| | 35 | is_disordered_040 | 1 if disorder_fraction is ≥ the 40th percentile of disorder fraction within DisProt for S288C proteins, otherwise 0 |
| | 36 | is_disordered_0.45 | 1 if disorder_fraction is ≥ the 45th percentile of disorder fraction within DisProt for S288C proteins, otherwise 0 |
| | 37 | is_disordered_0.50 | 1 if disorder_fraction is ≥ the 50th percentile of disorder fraction within DisProt for S288C proteins, otherwise 0 |
| | 38 | is_disordered_0.55 | 1 if disorder_fraction is ≥ the 55th percentile of disorder fraction within DisProt for S288C proteins, otherwise 0 |
| | 39 | is_disordered_0.60 | 1 if disorder_fraction is ≥ the 60th percentile of disorder fraction within DisProt for S288C proteins, otherwise 0 |
| | 40 | is_disordered_0.65 | 1 if disorder_fraction is ≥ the 65th percentile of disorder fraction within DisProt for S288C proteins, otherwise 0 |
| | 41 | is_disordered_0.70 | 1 if disorder_fraction is ≥ the 70th percentile of disorder fraction within DisProt for S288C proteins, otherwise 0 |
| | 42 | is_disordered_0.75 | 1 if disorder_fraction is ≥ the 75th percentile of disorder fraction within DisProt for S288C proteins, otherwise 0 |
| | 43 | is_disordered_0.80 | 1 if disorder_fraction is ≥ the 80th percentile of disorder fraction within DisProt for S288C proteins, otherwise 0 |
| | 44 | is_disordered_0.85 | 1 if disorder_fraction is ≥ the 85th percentile of disorder fraction within DisProt for S288C proteins, otherwise 0 |
| | 45 | is_disordered_0.90 | 1 if disorder_fraction is ≥ the 90th percentile of disorder fraction within DisProt for S288C proteins, otherwise 0 |
| | 46 | is_disordered_0.95 | 1 if disorder_fraction is ≥ the 95th percentile of disorder fraction within DisProt for S288C proteins, otherwise 0 |
| | 47 | is_disordered_1.00 | 1 if disorder_fraction is ≥ the 100th percentile of disorder fraction within DisProt for S288C proteins, otherwise 0 |
| | 48 | albatross | Dictionary containing predicted IDR conformational ensemble properties (keys are integers, matches IDR_sequences) |
| | 49 | cider | Dictonary containing IDR sequence properties (keys are integers, matches IDR_sequences) |

### Supplementary Table 1 (cont.) – Annotations within ANYI

| Annotation Class | Index | Data Column Name | Explanation |
|---|---|---|---|
| 5 - Predicted protein structure & stability | 50 | structure_sequence | Amino acid sequence from EBI AlphaFold2 structure as a string |
| | 51 | cleaved_structure_sequence | Sequence for this protein after removal of signal sequence |
| | 52 | sequence_matches_structure | TRUE if the sequence from the EBI AlphaFold2 structure matches the sequence from SGD, otherwise FALSE |
| | 53 | final_sequence | Final amino acid sequence matching the final structure |
| | 54 | final_structure_source | Source of the final structure selected from the list: {EBI, AF2-cleaved, AF2, None}. EBI = the original structure from the EBI AlphaFold2 database; AF2-cleaved = the original structure from the EBI with the signal sequence removed; AF2 = a new AlphaFold2 prediction; None = No structure available (typically due to a lack of a verified sequence) |
| | 55 | mean_plddt | Average of C-alpha pLDDT values for all proteins with a structure |
| | 56 | L | Length of the final_sequence for each protein |
| | 57 | Ghosh-Dill-dG | Free energy difference between folded and unfolded states for this protein calculated with Eq. 1 of Ghosh-Dill 2010 (kcal/mol) |
| | 58 | cagiada-dG | Absolute free energy difference between folded and unfolded states for this protein calculated with Cagiada language model (kcal/mol) |
| | 59 | Rosetta_total_score_0001 | Value of Rosetta total_score with the ref2015 scoring function for the first FastRelax replicate |
| | 60 | Rosetta_total_score_0002 | Value of Rosetta total_score with the ref2015 scoring function for the second FastRelax replicate |
| | 61 | Rosetta_total_score_0003 | Value of Rosetta total_score with the ref2015 scoring function for the third FastRelax replicate |
| | 62 | Rosetta_total_score_0004 | Value of Rosetta total_score with the ref2015 scoring function for the fourth FastRelax replicate |
| | 63 | Rosetta_total_score_0005 | Value of Rosetta total_score with the ref2015 scoring function for the fifth FastRelax replicate |
| | 64 | Rosetta_total_score_0006 | Value of Rosetta total_score with the ref2015 scoring function for the sixth FastRelax replicate |
| | 65 | Rosetta_total_score_0007 | Value of Rosetta total_score with the ref2015 scoring function for the seventh FastRelax replicate |
| | 66 | Rosetta_total_score_0008 | Value of Rosetta total_score with the ref2015 scoring function for the eighth FastRelax replicate |
| | 67 | Rosetta_total_score_0009 | Value of Rosetta total_score with the ref2015 scoring function for the ninth FastRelax replicate |
| | 68 | Rosetta_total_score_0010 | Value of Rosetta total_score with the ref2015 scoring function for the tenth FastRelax replicate |
| | 69 | Rosetta_best_total_score | Best (lowest) value of Rosetta total_score with the ref2015 scoring function across all 10 FastRelax replicates |
| | 70 | Rosetta_best_pose_index | Index of pose with best total_score |

### Supplementary Table 1 (cont.) – Annotations within ANYI

| Annotation Class | Index | Data Column Name | Description |
|---|---|---|---|
| 6 - Protein half life | 71 | Christiano_degradation_rate(min-1) | Degradation rate in inverse minutes from Christiano et al. 2014 |
| | 72 | Christiano_degradation_R2 | Pearson correlation coefficient from Christiano et al. 2014 curve fitting |
| | 73 | Christiano_halflife_min | Halflife in minutes from Christiano et al. 2014 |
| | 74 | Villen_halflife_hours | Halflife in hours from Martin-Perez and Villen 2017 |
| | 75 | Villen_halflife_SD_hours | Halflife standard deviation in hours from Martin-Perex and Villen 2017 |
| | 76 | Villen_halflife_CV | Halflife coefficient of variation from Martin-Perex and Villen 2017 |
| | 77 | Villen_halflife_min | Halflife in minutes from Martin-Perex and Villen 2017 |
| | 78 | Villen_halflife_SD_min | Halflife standard deviation in minutes from Martin-Perex and Villen 2017 |
| 7 - Protein expression level | 79 | mean_molecules_per_cell | Mean molecules per cell from Ho et al. 2018 |
| | 80 | median_molecules_per_cell | Median molecules per cell from Ho et al. 2018 |
| | 81 | expression_CV | Coefficient of variation of expression measurements from Ho et al. 2018 |
| 8 - Protein translation efficiency | 82 | log2_TE | log base 2 of translation efficiency from SciKit-ribo for Weinberg 2016 Ribo-Seq data |
| 9 - Post-translational modifications | 83 | ptm_bronze | Phosphorylation sites with "Bronze" evidence from PTMeXchange |
| | 84 | ptm_gold | Phosphorylation sites with "Gold" evidence from PTMeXchange |
| | 85 | ptm_silver | Phosphorylation sites with "Silver" evidence from PTMeXchange |
| | 86 | Acetylation (K) | List of lysine residue numbers that are predicted by PTMGPT2 to be acetylated |
| | 87 | Amidation (V) | List of valine residue numbers that are predicted by PTMGPT2 to be amidated |
| | 88 | Formylation (K) | List of lysine residue numbers that are predicted by PTMGPT2 to be formylated |
| | 89 | Glutarylation (K) | List of lysine residue numbers that are predicted by PTMGPT2 to be glutarylated |
| | 90 | Glutathionylation (C) | List of cysteine residue numbers that are predicted by PTMGPT2 to be glutathionylated |
| | 91 | Hydroxylation (K) | List of lysine residue numbers that are predicted by PTMGPT2 to be hydroxylated |
| | 92 | Hydroxylation (P) | List of proline residue numbers that are predicted by PTMGPT2 to be hydroxylated |
| | 93 | Malonylation (K) | List of lysine residue numbers that are predicted by PTMGPT2 to be malonylated |
| | 94 | Methylation (K) | List of lysine residue numbers that are predicted by PTMGPT2 to be methylated |
| | 95 | Methylation (R) | List of arginine residue numbers that are predicted by PTMGPT2 to be methylated |
| | 96 | N-linked Glycosylation (N) | List of asparagine residue numbers that are predicted by PTMGPT2 to be N-glycosylated |
| | 97 | O-linked Glycosylation (S,T) | List of serine or threonine residue numbers that are predicted by PTMGPT2 to be O-glycosylated |
| | 98 | Phosphorylation (S,T) | List of serine or threonine residue numbers that are predicted by PTMGPT2 to be phosphorylated |
| | 99 | Phosphorylation (Y) | List of tyrosine residue numbers that are predicted by PTMGPT2 to be phosphorylated |
| | 100 | S-nitrosylation (C) | List of cysteine residue numbers that are predicted by PTMGPT2 to be S-nitrosylated |
| | 101 | S-palmitoylation (C) | List of cysteine residue numbers that are predicted by PTMGPT2 to be S-palmitoylated |

### Supplementary Table 1 (cont.) – Annotations within ANYI

| Annotation Class | Index | Data Column Name | Description |
|---|---|---|---|
| 9 - Post-translational modifications | 102 | Succinylation (K) | List of lysine residue numbers that are predicted by PTMGPT2 to be succinylated |
| | 103 | Sumoylation (K) | List of lysine residue numbers that are predicted by PTMGPT2 to be sumoylated |
| | 104 | Ubiquitination (K) | List of lysine residue numbers that are predicted by PTMGPT2 to be ubiquitinated |
| 10 - Entanglement status | 105 | AF2_contains_entanglement | 1 if AlphaFold2 structure for this protein contains an entanglement, otherwise 0. If a protein did not have its entanglement assessed, its value will be 0 |
| | 106 | exp_contains_entanglement | 1 if experimental structure for this protein contains an entanglement, otherwise 0. If a protein did not have its entanglement assessed, its value will be 0 |
| 11 - Chaperone interactions | 107 | interacting_chaperones | List of chaperones/co-chaperones with which this protein interacts |
| | 108 | interacting_chaperone_info | Additional information on the chaperone family, etc. |
| 12 - Oligomerization state | 109 | homodimer | 1 if protein is reported in Complex Portal to form a homodimer, otherwise 0 |
| | 110 | homotrimer | 1 if protein is reported in Complex Portal to form a homotrimer, otherwise 0 |
| | 111 | homotetramer | 1 if protein is reported in Complex Portal to form a homotetramer, otherwise 0 |
| | 112 | heterodimer | 1 if protein is reported in Complex Portal to form a heterodimer, otherwise 0 |
| | 113 | heterotrimer | 1 if protein is reported in Complex Portal to form a heterotrimer, otherwise 0 |
| | 114 | heterotetramer | 1 if protein is reported in Complex Portal to form a heterotetramer, otherwise 0 |
| | 115 | heteropentamer | 1 if protein is reported in Complex Portal to form a heteropentamer, otherwise 0 |
| | 116 | heterohexamer | 1 if protein is reported in Complex Portal to form a heterohexamer, otherwise 0 |
| | 117 | heteroheptamer | 1 if protein is reported in Complex Portal to form a heteroheptamer, otherwise 0 |
| | 118 | heterooctamer | 1 if protein is reported in Complex Portal to form a heterooctamer, otherwise 0 |
| | 119 | heterononamer | 1 if protein is reported in Complex Portal to form a heterononamer, otherwise 0 |
| | 120 | heterodecamer | 1 if protein is reported in Complex Portal to form a heterodecamer, otherwise 0 |
| | 121 | in_complex | 1 if protein is reported in Complex Portal to participate in formation of at least 1 complex, otherwise 0 |
| | 122 | complex_partners | List of other proteins that form stable complexes with this node |
| | 123 | complex_count_unknown | Integer number of complexes with unknown stoichiometry in which this protein is found |
| | 124 | complex_count | Integer number of complexes of known or unknown stoichiometry in which this protein is found |
| 13 - Domain annotations | 125 | PROSITE_PROFILE_Ndomains | Number of PROSITE PROFILE domains from InterPro for this protein |
| | 126 | PROSITE_PROFILE_domains | PROSITE PROFILE domain ranges for this protein |
| | 127 | Gene3D_Ndomains | Number of Gene3DD domains from InterPro for this protein |
| | 128 | Gene3D_domains | Gene3D domain ranges for this protein |
| | 129 | PANTHER_Ndomains | Number of PANTHER domains from InterPro for this protein |
| | 130 | PANTHER_domains | PANTHER domain ranges for this protein |
| | 131 | Pfam_Ndomains | Number of Pfam domains from InterPro for this protein |

### Supplementary Table 1 (cont.) – Annotations within ANYI

| Annotation Class | Index | Data Column Name | Description |
|---|---|---|---|
| | 132 | Pfam_domains | Pfam domain ranges for this protein |
| | 133 | SUPERFAMILY_Ndomains | Number of superfamily domains from InterPro for this protein |
| | 134 | SUPERFAMILY_domains | Superfamily domain ranges for this protein |
| | 135 | PROSITE_PATTERN_Ndomains | Number of PROSITE PATTERN domains from InterPro for this protein |
| | 136 | PROSITE_PATTERN_domains | PROSITE PATTERN domain ranges for this protein |
| | 137 | SMART_Ndomains | Number of SMART domains from InterPro for this protein |
| | 138 | SMART_domains | SMART domain ranges for this protein |
| | 139 | TIGRFAM_Ndomains | Number of TIGRFAM domains from InterPro for this protein |
| | 140 | TIGRFAM_domains | TIGRFAM domain ranges for this protein |
| 14 - Essentiality | 141 | essential | 1 if this protein is essential, 0 if it is not (from SGD function browser) |
| | 142 | essentiality_details | Detail of the essentiality finding |
| 15 - Yeast two-hybrid network data | 143 | Y2H_wkshell | Weighted k-shell decomposition result; computed with Cytoscape app wk-shell-decomposition |
| | 144 | Y2H_degree_centrality | Degree centrality; computed with NetworkX |
| | 145 | Y2H_betweenness_centrality | Betweenness centrality; computed with NetworkX |
| | 146 | Y2H_eigenvector_centrality | Eigenvector centrality; computed with NetworkX |
| | 147 | Y2H_closeness_centrality | Closeness centrality; computed with NetworkX |
| | 148 | Y2H_load_centrality | Load centrality; computed with NetworkX |
| | 149 | Y2H_pagerank | PageRank; computed with NetworkX with damping parameter = 0.85 |
| | 150 | Y2H_information_centrality | Information centrality; computed with NetworkX on the largest connected subgraph |
| 16 - Thermal proteome profiling | 151 | meltome-melting-point | Meltome Atlas melting temperature in degrees Celsius |
| | 152 | meltome-stability-class | Meltome Atlas stability class for this protein from the list {non-melter, stable, medium, non-stable} |
| | 153 | meltome-AUC | Area under the melting curve for this protein in Meltome Atlas |
| | 154 | meltome-fit-converged | True if the Meltome Atlas fit converged for this protein, otherwise np.nan |
| | 155 | meltome-fit-R2 | Pearson correlation coefficient for melting curve fit for this protein |
